# Parent attention-orienting behavior is associated with neural entropy in infancy

**DOI:** 10.1101/2024.03.14.585061

**Authors:** Cabell L. Williams, Allison R. Belkowitz, Madelyn G. Nance, Emily T. Mortman, Soni Bae, Sheher-Bano Ahmed, Meghan H. Puglia

**Affiliations:** University of Virginia, Department of Psychology; University of Virginia, Department of Neurology; Smith College, Department of Neuroscience

**Keywords:** multiscale entropy, infant development, joint attention, EEG, parent behavior

## Abstract

Parents play a significant role in directing infant’s attention to environmental stimuli via joint attention. We hypothesized that infants whose parents provide more bids for joint attention will display a more complex neural response when viewing social scenes. Sixty-one 8-month-old infants underwent electroencephalography (EEG) while viewing videos of joint-and parallel-play and participated in a parent-infant free play interaction. EEG data was analyzed using multiscale entropy, which quantifies moment-to-moment neural variability. Free play interactions were coded for parent alternating gaze, a behavioral mechanism for directing attention to environmental cues. We found a significant positive association between parent alternating gaze and neural entropy in frontal and central brain regions. These results suggest a relationship between parent behavior and infant neural mechanisms that regulate social attention, underlying the importance of parent cues in the formation of neural networks in infancy.

## Background

The early social environment primarily consists of interactions between the infant and caregiver. Caregiver social cues enable the infant to engage with their environment (Moore & Dunham, 1995; Tomasello, 1995). Joint attention is a preverbal referential behavior by which individuals follow or direct another’s attention (Bruner, 1974) that emerges around six months (Butterworth & Jarrett, 1991). Joint attention between caregivers and infants has been associated with consolidation of social information (Mundy & Gomes, 1998), language acquisition (Mundy & Gomes, 1998; Salo et al., 2018; Tomasello & Farrar, 1986), greater visual attention to their partner (Striano & Stahl, 2005), proficiency in culturally normative behavior (Bruner, 1974), and educational success (Mundy & Newell, 2007). A key parent-directed joint attention behavior occurs when parents alternate gaze between the infant and an object (Bakeman & Adamson, 1984). The effects of parent-, rather than infant-directed joint attention (e.g., using gestures to direct caregiver attention (Carpenter et al., 1998)), on infant neural responses to social information has not yet been explored. This research aims to assess how early caregiving environment impacts infant social-attentional brain function.

Infancy is a sensitive period of development characterized by extensive neural plasticity. Infant brains are highly susceptible to neural rewiring based on experience (Bennett et al., 2018; Kolb et al., 2017). Neural development results in greater functional variability and neurological complexity (McIntosh et al., 2010). Neural entropy – a measure of functional variability and complexity – has been associated with the exchange of information between neurons (Shew et al., 2009, 2011) and increased neural synchrony (Mišić et al., 2015), and may facilitate the orientation to and detection of important environmental signals. Here, we test the hypothesis that infants whose parents provide a more socially-oriented environment (i.e., more joint attention bids through parent alternating gaze) will demonstrate increased neural entropy in regions supporting attention orienting (Hopfinger et al., 2000) and social perception (Grossmann, 2015) when viewing children engaging in joint-play compared to parallel-play.

## Methods

### Participants

Sixty-nine infants were recruited from the University of Virginia’s (UVA) hospital database and the greater Charlottesville area as part of a longitudinal study at birth, 4, 8, 12 and 16 months. Given that joint attention emerges around 6 months (Butterworth & Jarrett, 1991), we assessed neural and behavioral data at 8 months (*M*=248.56 days, *SD*=13.22), as this was the earliest timepoint by which most of our participants likely developed joint attention abilities.

Infants completed a free play interaction with their parent and underwent EEG. The parent provided written informed consent as approved by UVA’s Institutional Review Board, and was paid $50 for each visit. Eight participants were excluded due to insufficient artifact-free EEG data (see EEG Acquisition and Preprocessing). The final dataset consisted of 61 (33 female) infants.

### Infant Behavior

The parent and infant underwent a 5-minute videotaped naturalistic free play interaction in which they were instructed to play as they would at home. Videos were later coded by a research assistant blind to the hypothesis. The first minute of each video was discarded to allow the dyad to acclimate to the environment. The videos were coded for duration of parent alternating gaze, defined as the parent directing visual attention to their infant, then to an object, and back again for maximum 2-s per turn. Videos were coded using INTERACT (Mangold, Germany). Twenty-five percent of randomly-selected videos were coded by a secondary coder to establish interrater reliability (two-way random effects model, r=.81) (Koo & Li, 2016).

### EEG Acquisition and Preprocessing

Infants viewed 16-s videos of two children engaging in joint- or parallel-play while undergoing EEG. Video order was pseudorandomized such that no child pair was shown back-to-back, and no play viewing condition (joint/parallel) repeated more than twice. The next video began once the infant fixated on an attention-getter at screen center for 500 ms, as measured by a Pro Spectrum (Tobii, Sweden) eye tracker. The videos continued until the infant failed to orient to the screen for six consecutive trials or all 16 trials were completed. Visual attention to the paradigm was assessed via eye tracking, video recording, and experimenter observation. The videos were presented using PsychToolBox v3.0.14 (Brainard, 1997) for MATLAB (Mathworks, Natick, MA). Infants were seated on their parent’s lap during EEG collection. Parents were instructed not to interact with the infant until completion of the paradigm and wore sunglasses to prevent the eye tracker from registering their gaze.

EEG data was collected using 32 Ag/AgCl active actiCAP slim electrodes (Brain Products GmbH, Germany) fastened to an elastic cap following the 10-20 electrode placement system (Figure 1). EEG signals were amplified using a BrainAmp DC Amplifier and recorded with BrainVision Recorder software at a sampling rate of 5000 Hz with an online reference to FCz, and an online band-pass filter between 0.01-1000 Hz.

**Figure 1.**
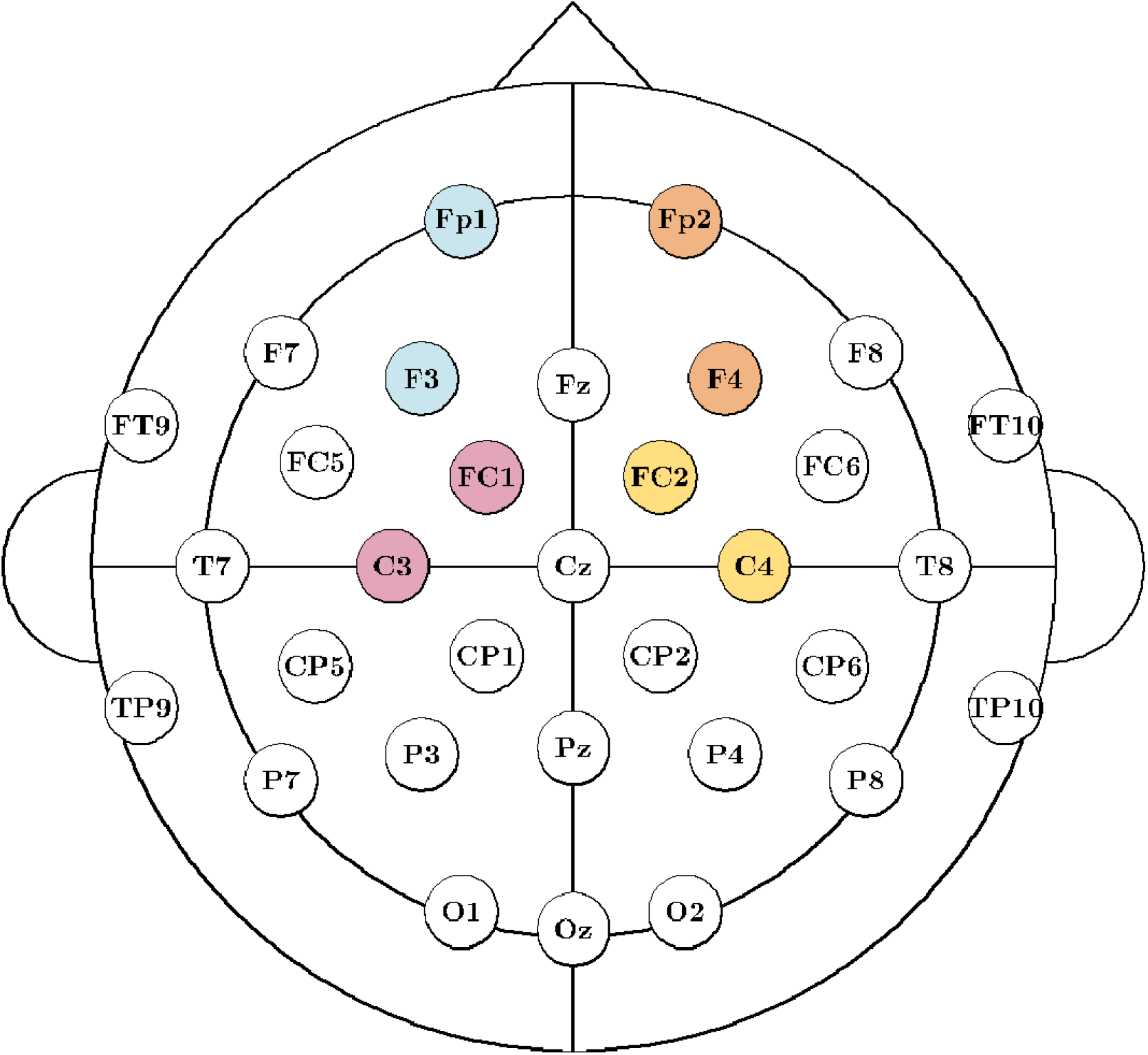
Infant EEG electrode montage. EEG data from 3-8 Hz was averaged together to create the following regions of interest: Left frontal: Fp1 and F3 (blue); Right frontal Fp2 and F4 (orange); Left central: FC1 and C3 (red); Right central: FC2 and C4 (yellow). All other scalp electrodes were grouped as a comparison region of interest.

EEG preprocessing was completed using the Automated Preprocessing Pipe-Line for the Estimation of Scale-wise Entropy from EEG Data (APPLESEED) (Puglia et al., 2022), a standardized EEG preprocessing and entropy estimation pipeline for pediatric EEG data implemented through EEGLab (Delorme & Makeig, 2004). We first downsampled the data to 500 Hz, applied a 0.1-100 Hz bandpass filter, and removed 60 Hz sinusoidal noise using CleanLine (2021). Sixteen 1-s epochs were extracted from each video. Channels that exceeded 500 μV in greater than 50% of epochs were spherically interpolated. Epochs contaminated with excessive amplitude standard deviations (>80 μV) within a 200-ms sliding window were discarded. The data was re-referenced to the average of all scalp electrodes. Because the number of data points influences the reliability of multiscale entropy (MSE) estimates (Grandy et al., 2016), it is critical that MSE is computed from an equivalent number of data points for each participant. Therefore, the 10 trials for each viewing condition with a total global field power (GFP) closest to the median GFP for each participant were selected for downstream analysis. Ten trials per condition is the most common threshold employed in infant EEG studies (Monroy et al., 2021) and allowed us to retain the maximum number of participants. Eight participants had fewer than 10 artifact-free trials in each condition and were excluded from subsequent analysis. The final dataset consisted of 61 infants.

### Analysis

MSE quantifies signal variability and irregularity across time scales (Costa et al., 2002) by measuring iterative patterns within a time series and assigning low values to predictable patterns and high values to unpredictable patterns. Neural systems operate on multiple timescales simultaneously, necessitating the use of a multiscale approach for measuring neural signal variability. MSE analysis was computed using APPLESEED (Puglia et al., 2022) with a pattern length *m* = 2 and similarity criterion *r* = 0.5. Critically, this pipeline uses a modified MSE algorithm that recomputes *r* at each scale (Nikulin & Brismar, 2004) and computes MSE across discontinuous segments (Grandy et al., 2016) enabling entropy estimates at more fine-grained time scales. Entropy estimates within the 3 to 8 Hz range – the predominant frequency in 8-month-old infants (Saby & Marshall, 2012) – were averaged together within four regions of interest (ROIs): left frontal (Fp1, F3), right frontal (Fp2, F4), left central (FC1, C3), and right central (FC2, C4) – selected as regions implicated in attention orienting (Hopfinger et al., 2000) and social perception (Grossmann, 2015). All other electrodes were grouped as a comparison ROI (Figure 1).

Statistics were run in RStudio v2022.12.0 (RStudio Team, 2020). A general linear model assessed the association between parent alternating gaze and MSE with viewing condition (joint vs. parallel) and ROI (left frontal, right frontal, left central right central, other) included as covariates. Parent alternating gaze was *z*-score transformed; the square root of MSE was calculated to validate the assumption of normally distributed residuals. Outlier data points were identified as those for which the absolute value of the median standardized residuals was >3. Three outliers in the joint-play condition (1 left frontal, 1 right frontal, and 1 right central) were removed. Then, we tested for data points with undue influence on the model using Cook’s distance (D) > 1 (Cook & Weisberg, 1982) as criteria. No influential points (all Ds < 0.02) were detected. Post-hoc comparisons of estimated marginal means were used to explore differences across investigated ROIs via the emmeans R package (Lenth, 2018).

## Results

We found a significant positive effect of parent alternating gaze (β=.013, p=.042) on entropy, but no effect of viewing condition (β=.007, p=.512) or ROI (all ps >.361). While post-hoc comparisons revealed no statistically significant differences between ROIs (all ps > .636), a visual examination of the plotted associations revealed trending associations among our considered variables (Figure 2). Specifically, within the joint-play viewing condition, we identified a trending positive association between parent alternating gaze and infant neural entropy in frontal regions, whereas we identified a trending negative association between these variables in central and other brain regions. However, during parallel-play viewing, parent alternating gaze was positively associated with infant neural entropy in all brain regions.

**Figure 2.**
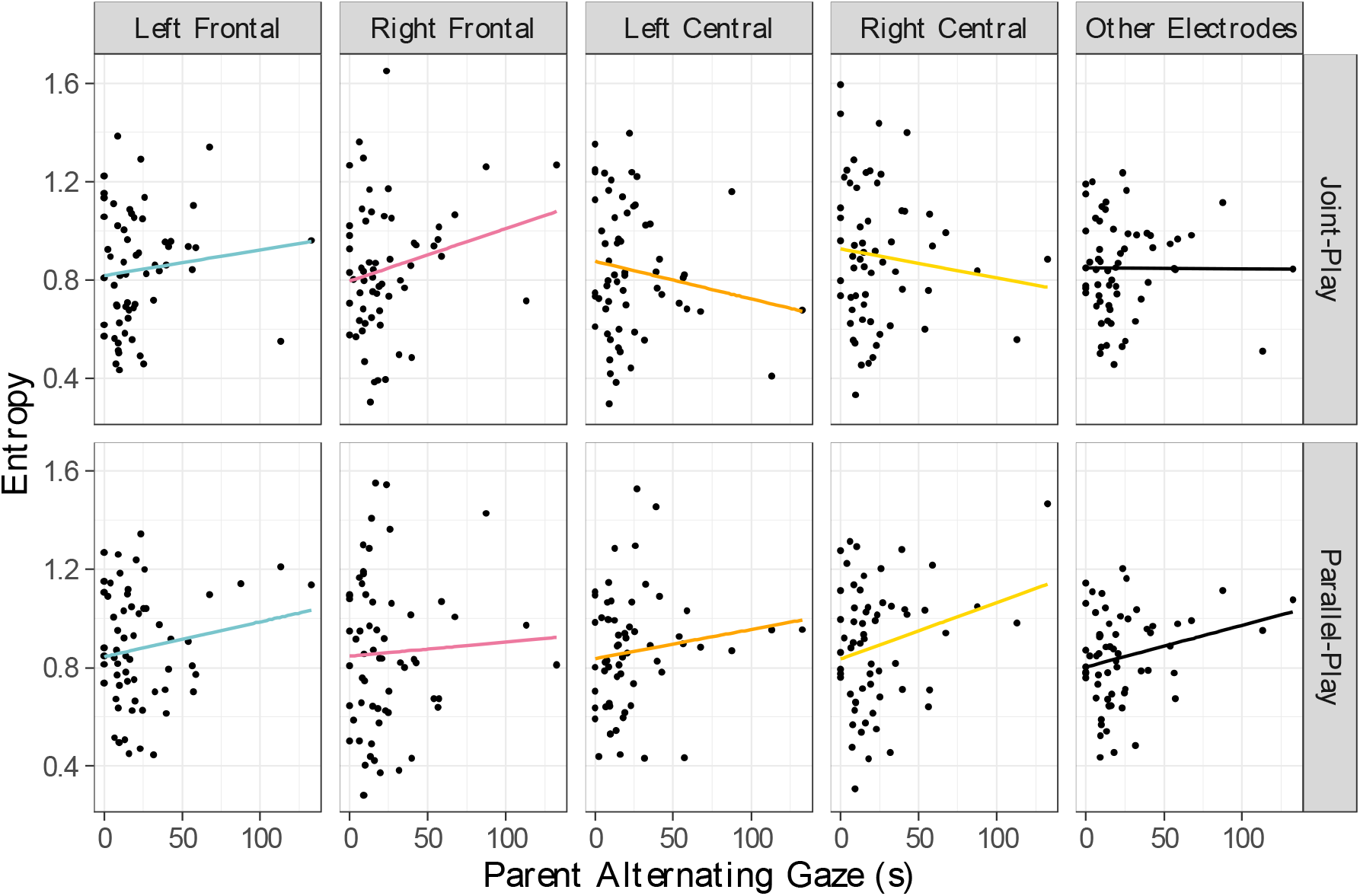
Association between parent alternating gaze and multiscale entropy. Multiscale entropy (MSE) was computed on infant EEG data from 3-8 Hz for five regions of interest (vertical panels) over two play viewing conditions (horizontal panels). Parent alternating gaze is significantly associated with MSE (β=.013, p=-.042). While there is no significant effect of play condition (β=.0087 p=.512) or region on interest (all ps >.361), there is a trending shift from a positive association in the frontal lobe (blue, red) to a negative association in the central lobe (orange, yellow) specific to the social processing (joint play viewing) condition (top panel).

## Discussion

This study assessed parent gaze orienting as an important environmental factor in the modulation of infant social attention through increased neural variability. Our results showed a positive association between parent alternating gaze and neural entropy in 8-month-old infants. At 8 months, infants are developing attentional control (Stroganova et al., 1998), a process facilitated by parent gaze cues which helps infants regulate attention to relevant external stimuli (Salley et al., 2016). This skill enables increased mutual gaze between infants and their parents, which is key for developing aspects of sociality (e.g., sense of self and regulation of emotion) (Niedźwiecka et al., 2018).

Our finding that parent alternating gaze is associated with increased neural entropy aligns with current literature surrounding the functionality of signal variability. Increased entropy represents a more complex, mature system capable of exploring varying brain states (Wang et al., 2018). A positive association between parent gaze cues and neural entropy infers that the infant finds social information maximally salient (Puglia et al., 2018) and may reflect accelerated development (Paterson et al., 2006).

We found trending effects of viewing condition (joint-vs. parallel-play) on the association between MSE and parent behavior across central and frontal cortices. The association trends positive in frontal and negative in central regions for the joint-play viewing condition only, suggesting that 8-month-old infants discriminate social interactions within brain regions specifically involved with attention regulation (Hopfinger et al., 2000) and social information processing (Grossmann, 2015).

While significant, the effect sizes and variance explained was low. Future investigations should replicate these results with a larger sample, consider additional social environmental influences, and longitudinally assess associations between parent behavior and infant social-attentional development (Cleveland & Striano, 2007).

### Conclusions

Neural plasticity occurs as environmental factors shape an infant’s brain development. Parent gaze is an important environmental stimulus that facilitates an infant’s ability to perceive and interpret external stimuli, which is critical to the development of neural networks that underlie social skills. We demonstrated that parent-directed social orienting cues modulate neural variability reflective of attentional states during social processing. This research highlights the importance of early caregiving on emerging social attentional skills in infancy.

## Acknowledgements

This work was supported by an NSF EXPAND fellowship to CLW, the National Institute of Mental Health K01MH125173 to MHP, and the Jefferson Trust Foundation to MHP. We thank participating families for taking part in our research. The authors declare no conflicts of interest.

## Notes

The data that support the findings of this study are available from the corresponding author upon reasonable request.

### Competing Interest Statement

The authors have declared no competing interest.

